# Increased random exploration in schizophrenia is associated with inflammation

**DOI:** 10.1101/2020.03.15.989483

**Authors:** Flurin Cathomas, Federica Klaus, Karoline Guetter, Hui-Kuan Chung, Anjali Raja Beharelle, Tobias Spiller, Rebecca Schlegel, Erich Seifritz, Matthias N. Hartmann-Riemer, Philippe N. Tobler, Stefan Kaiser

**Author notes:** Department of Psychiatry, Psychotherapy and Psychosomatics, Psychiatric Hospital, University of Zurich, Lenggstrasse 31, 8032 Zürich, Switzerland (Flurin Cathomas). Division of Adult Psychiatry, Department of Psychiatry, Geneva University Hospitals, Chemin du Petit-Bel-Air, 1225 Chêne-Bourg, Switzerland (Stefan Kaiser).

## Abstract

One aspect of goal-directed behavior, which is known to be impaired in patients with schizophrenia (SZ), is balancing between exploiting a familiar choice with known reward value and exploring a lesser known but potentially more rewarding option. Despite its relevance to several symptom domains of SZ, this has received little attention in SZ research. In addition, while there is increasing evidence that SZ is associated with chronic low-grade inflammation, few studies have investigated how this relates to specific behaviors, such as balancing exploration and exploitation. We therefore assessed behaviors underlying the exploration-exploitation trade-off using a three-armed bandit task in 45 patients with SZ and 19 healthy controls (HC). This task allowed us to dissociate goal-unrelated (random) from goal-related (directed) exploration and correlate them with psychopathological symptoms. Moreover, we assessed a broad range of inflammatory proteins in the blood and related them to bandit task behavior. We found that, compared to HC, patients with SZ showed reduced task performance. This impairment was due to a shift from exploitation to random exploration, which was associated with symptoms of disorganization. Relative to HC, patients with SZ showed a pro-inflammatory blood profile. Furthermore, high sensitivity C-reactive protein (CRP) positively correlated with random exploration, but not with directed exploration or exploitation. In conclusion, we show that low-grade inflammation in patients with SZ is associated with random exploration, which can be considered a behavioral marker for disorganization. CRP may constitute a marker for severity of, and a potential treatment target for maladaptive exploratory behaviors.

## Introduction

Schizophrenia is a complex neuropsychiatric disorder characterized by a wide range of symptoms [1]. Negative symptoms and disorganization are strongly associated with reduced social and occupational functioning and are challenging to treat [2] [3] [4] [5]. Therefore, understanding the shared and dissociable mechanisms underlying these symptoms is crucial in increasing our knowledge about the etio-pathophysiology of the disorder and the much needed development of novel treatments.

Negative symptoms and disorganization have been linked to impairments of goal-directed behaviors [6] [7]. Investigating behaviors relevant to these symptom dimensions in schizophrenia with objective, task-based measures that allow the assessment of complex, well defined behaviors with ecological validity is crucial [8] [9]. The exploration-exploitation trade-off concerns the decision between exploiting a familiar option with known reward value and exploring a lesser known option with more uncertain but potentially higher reward value. This trade-off may constitute a paradigmatic case for assessing goal-directed behaviors, which have been shown to be impaired in patients with schizophrenia [10] [11]. To study the exploration-exploitation trade-off in a controlled experimental setting, a human computerized task, where the participant has to repeatedly choose between different “slot-machines” (bandits) that gradually change their reward values has been developed [12] [13]. Accordingly, participants have to decide when to switch from exploitative to exploratory strategies when aiming to obtain the highest total reward pay-off possible [13] [14]. An optimal balance between exploration and exploitation results in a maximization of the obtained reward and fulfils the fundamental need of foraging organisms to adapt to a complex and changing world (for review [15]).

Few studies have applied exploration-exploitation tasks in patients with schizophrenia (SZ), but no firm conclusions can be drawn so far: Strauss and colleagues found that patients with SZ show decreased uncertainty-driven directed exploration, which was associated with clinically assessed negative symptoms [16]. Another study reported that patients with SZ displayed increased novelty-seeking behaviors [17]. In addition, while several brain regions (midbrain and prefrontal areas) and neurotransmitters (dopamine, noradrenaline and acetylcholine) have been implicated in mediating exploration-exploitation trade-offs [10] [15] [18] [19], the underlying etiological mechanisms remain to be elucidated.

Inflammation has been suggested to be a key mechanism for disturbances of motivation and goal-directed behavior [20] [21]. Pre-clinical animal models and human studies have shown a close interaction between the peripheral immune system and the central nervous system in health and the development of neuropsychiatric disorders [22] [23]. Indeed, several studies have reported that patients with schizophrenia display alterations in immune function indexed by cytokines, chemokines and acute phase proteins in both blood and cerebral spinal fluid (CSF), which are indicative of chronic low-grade inflammation (see [24] for meta-analysis).

Recent studies have started to investigate whether these immune changes correlate with specific behaviors [25]. With regard to exploration and exploitation, the concept of sickness behavior as an adaptive behavioral response to acute infections is of high relevance [26] [27]. Sickness behavior can be interpreted as a short-term disturbance of exploratory behaviors that could have beneficial energy-saving effects by shifting energy resources towards the immune system to facilitate clearing infections and wound healing. However, in our modern world with fewer infectious challenges and better sanitation, chronic-low grade inflammation might be maladaptive leading to inefficient performance [28]. To our knowledge, no study has so far investigated the relationship of a pro-inflammatory state with an objective task assessment of exploration-exploitation behavior, despite the high relevance for several symptom domains of SZ. Overall, we hypothesized that patients with SZ will display reduced task outcome based on impairments in balancing between exploration and exploitation behaviors and that the underlying behavioral impairments are associated with a low-grade, pro-inflammatory state.

Therefore, we first assessed negative, disorganized and cognitive symptoms in pharmacologically stable patients with SZ and healthy controls (HC). We then examined exploration vs. exploitation behaviors using a three-armed computerized bandit task and investigated correlations between overall task performance, directed and random exploration and exploitation with psychopathological measures. Next, we assessed group differences (SZ vs. HC) in a broad array of cytokines and chemokines in the blood. To investigate and visualize the complex interaction between the individual inflammatory parameters and their relationship with group affiliation (SZ or HC) we used network analysis. Lastly, we investigated the relationships between the immune-markers that were different between groups and task-based behaviors.

## Methods

### 2.1. Participants

45 patients meeting the DSM-V (American Psychiatric Association, 2013) criteria for SZ and 19 HC subjects were recruited. Patients were recruited from outpatient and inpatient units of the Psychiatric Hospital of the University of Zurich and affiliated institutions, HC were recruited from the community via advertisement. More patients than controls were recruited to have adequate power for the correlational analyses (see below). Two patients did not complete the bandit task (due to fatigue) and are therefore only included in the group comparison of immune markers. Diagnoses were confirmed by conducting the Mini-International Neuropsychiatric Interview [29]. All patients were clinically stable and under a stable dose of medication for at least two weeks prior to testing. Inpatients were at the end of their hospitalization and engaged in a multimodal therapy program and activities outside the hospital. The average duration of hospitalization for patients with schizophrenia in Swiss psychiatric hospitals is longer than in most other countries, so the majority of patients would have been treated as outpatients in other health care systems. The inclusion age was between 18-65 years. We excluded patients with any other than the above mentioned DSM-IV Axis I disorders, lorazepam medication higher than 1mg per day and acute psychotic symptoms. Participants with any alcohol use disorder based on lifetime criteria and participants with current cannabis abuse or dependency or with any other substance abuse were excluded. HC were excluded if any neuropsychiatric diagnosis was present in the structured Mini-International Neuropsychiatric Interview. In both groups, participants were excluded if they had a history of head-injury or any autoimmune or chronic inflammatory disorder or if they took any pain-medication or anti-inflammatory drugs at least one week prior to testing (assessed by detailed questionnaire and medical records where available). Furthermore, participants were not included in the study if they had a history of any known acute inflammation two weeks prior to testing. All participants gave written informed consent and the project was approved by the Ethics Committee of the Canton of Zurich.

### 2.2. Assessment of Psychopathology and Cognition

Negative, positive and disorganized symptoms were assessed using the five factor model of the Positive and Negative Syndrome Scale (PANSS) [30]. Cognition was assessed with the Brief Neurocognitive Assessment (BNA) [31], which was shown to be highly correlated with the MATRICS Consensus Cognitive Battery (MCCB) [32] and has similarly good validity criteria [33]. With the BNA, a cognitive score is computed for each participant by combining results from the Letter-Number-Span Test and the Symbol Coding Test. Global level of functioning was assessed using the Personal and Social Performance Scale (PSP) [34].

### 2.3. Bandit Task

For assessing exploratory versus exploitative behavior, the participants performed a computerized virtual slot machine (bandit) task (**Fig. S1**). The procedure is an adapted version of a recently described bandit task [14]. Briefly, participants were shown three virtual slot machines depicted by 3 blue boxes on a screen between which they had to choose repeatedly by pressing the arrow keys on the keyboard. After each choice, the points they won, ranging from 1-100, were displayed. Participants were instructed to collect as many points as possible to maximize their gain. In foraging situations, like in this bandit task, decision makers need to balance exploitation with exploration. They were informed that the points paid out by the slot machines oscillated randomly and independently over trials, so they would have to choose at any given point between exploiting the currently used bandit and exploring the other two options to maximize gain. Unlike standard slots, the mean payoffs changed randomly and independently from trial to trial, with subjects finding information about the current worth of a slot only through sampling it [13]. Three brief written questions and a short practice trial to confirm the comprehension of the task took place before the actual task began. The payouts for each bandit were generated with a decaying Gaussian random walk, resulting in points paid off varying noisily around three different means. Reward of the selected option was displayed on the computer screen after each choice. If participants took too much time (5 seconds) deciding between the slot machines, a window popped up asking them to choose a machine. Participants completed 200 trials. At the end, five trials were randomly selected and the participants received the mean of the corresponding gains. To test whether participants performed better than chance, we simulated random choosers (equal chance to select each bandit) 1000 times for each participant’s reward structure. The average total reward earned by the random choosers was 51.90 points. We tested whether control participants and participants with SZ performed better or worse than the average of random choosers. To avoid any unnecessary assumptions about the actual implementation of psychological functions, we performed model-free analyses. We defined the best, second-best and worst options as the options with the highest, second highest and lowest expected value in the current trial based on the objective reward structure of the task. Continuously selecting the best option was defined as exploitation. There was a significant correlation between the average length of consecutively selecting the best option and points earned in the task (r(s) = 0.53, *p* < 0.001), further validating our measure of exploitation. Since the points paid out by the slot machines oscillated randomly and independently over trials, exploitative behavior will result in nonoptimal condition where people have insufficient information about the current reward structure. Ideally, decision makers should balance exploitation with exploration. To further disentangle different forms of exploratory behaviors, we distinguished directed from random exploration based on the uncertainty associated with each option [35] [36] [37]. The less often a given bandit has been chosen, the less information the decision maker has about the bandit. Accordingly, *directed exploration* corresponds to choosing the option that provides most information for future decisions, and can be viewed as goal-directed uncertainty reduction. In contrast, *random exploration* can be interpreted as a noisy behavior, which provides little or no extra information. In our task, directed exploration corresponds to a shift to the least selected option while shifts to more often selected options correspond to random exploration. A less-well established alternative definition of directed and random exploration is based on the value of each option, with value-based directed exploration defined as switching from the best option to the second best option. In the supplemental material, we show that the main findings of the present study remained qualitatively the same using this alternative definition (**Fig. S2**).

### 2.4. Blood samples and processing

Blood was drawn between 8-10 am on the day of testing. Study participants were instructed to fast for at least 8 hours prior to the blood draw and this was verified by a questionnaire on the day of testing. To obtain plasma for Olink panel, blood was drawn into ethylenediaminetetraacetic acid (EDTA) tubes (Sarstedt, Switzerland), centrifuged for 15min at 1500g and plasma was frozen at −80°C. High sensitivity C-reactive protein (CRP) was measured in serum: blood was collected into a silica and gel containing tube (BD Vacutainer).

### 2.5. CRP

CRP was measured in participant serum samples by immunoturbidimetry on Abbott Architect c16000 or c8000, at Unilabs Medical Analytics in Duebendorf / Zurich, Switzerland.

### 2.6. Cytokines and chemokines

Proteins were measured using the Olink® Inflammation panel (Olink Proteomics AB, Uppsala, Sweden) according to the manufacturer’s instructions. The Proximity Extension Assay (PEA) technology used for the Olink protocol is described in detail in [38]. In brief, pairs of oligonucleotide-labeled antibody probes bind to their targeted protein, and if the two probes are brought in close proximity the oligonucleotides will hybridize in a pair-wise manner. The addition of a DNA polymerase leads to a proximity-dependent DNA polymerization event, generating a unique PCR target sequence. The resulting DNA sequence is subsequently detected and quantified using a microfluidic real-time PCR instrument (Biomark HD, Fluidigm). Data is then quality-controlled and normalized using an internal extension control and an inter-plate control, to adjust for intra- and inter-run variation. The final assay readout is presented in Normalized Protein expression (NPX) values, which is an arbitrary unit on a log2-scale where a high value corresponds to a higher protein expression. Detection limits are available on the manufacturer’s website (www.olink.com). In total, 91 proteins were measured with an intra- and inter-assay coefficient of variance of < 10%. We excluded 19 proteins due to >80% values below the limit of detection (LOD). Values below the LOD were replaced with the LOD for the specific assay. 5 participants (2 HC, 3 SZ) were excluded because their samples failed quality control.

### 2.7. Data analysis

All variables were tested for normal distribution using the Kolmogorov-Smirnov test. If variables were normally distributed, potential group differences were examined using two-sided Students t-tests. To investigate correlations between inflammatory parameters and task variables / psychopathology, Pearson correlation coefficients r(p) were calculated. If data were not normally distributed, we instead used Mann Whitney U tests and Spearman correlation coefficients r(s). Heatmaps of inflammatory markers are based on standardized scores that were calculated as: (value of SZ (individual sample value) – mean of HCs)/ standard deviation of HCs). The level of significance was set at *p* < 0.05. To investigate group differences in inflammatory markers between HC and patients with SZ, we used an exploratory approach using uncorrected test statistics and network analysis (see below). Correlations between immune-markers and task behavior was controlled for multiple comparisons using the procedure proposed by Benjamini and Hochberg [39]. General linear models were used to test for the effects of covariates on group differences and partial correlations were calculated when controlling for covariates in the correlational analyses. Effect size was assessed by calculating Cohen’s d. Statistical analyses were computed with SPSS version 25 (IBM Corp., SPSS Inc., Chicago IL, USA), Stata version 14.2 (StataCorp. 2015. Stata Statistical Software: Release 14. College Station, TX: StataCorp LP), Matlab version R2018a (The MathWorks, Inc., Natick, Massachusetts, United States), R (Version 3.4.0, [40] and GraphPad Prism software (GraphPad Software Inc.). Heat maps were created using the online tool Morpheus (https://software.broadinstitute.org/morpheus).

### 2.8. Network analysis

Network analyses followed a four-step procedure: i) Data preparation: All variables for which the t-test for differences in blood levels of investigated inflammation markers between patients with SZ and HC had a *p*-value < 0.05 before correction for multiple testing were included in the network. ii) Network estimation and visualization: The estimated network is a Gaussian Graphical Model, in which the nodes represent the included variables. Edges between these variables can be interpreted as undirected, weighted partial correlations. Partial correlations can be thought of as the “unique” correlation between two nodes after controlling for the correlation with all other nodes [41]. We then visualized the network, with the thickness of an edge corresponding to its weight and with the color indicating a positive (red) or negative (blue) partial correlation between two nodes. Placement of the nodes for the visualization was based on the Fruchterman-Reingold-algorithm [42]. iii) Network characterization: We estimated centrality indices, which are a relative marker for the influence of a node on the network as a whole (for detail see: [43]). iv) Network accuracy and stability: Using current state of the art procedures, we calculated 95% confidence intervals for the edge weights and difference test for centrality and edge weights. All network analyses were carried out in R (Version 3.6.1, [40]) using the package bootnet [41]. Networks were visualized using qgraph [44]. Detailed information about network analysis are provided in the SI.

## Results

### 3.1. Sociodemographic data

Sociodemographic and clinical data of the sample are summarized in **Table 1**. Our SZ sample showed a wide range of negative and disorganized symptoms, and few positive symptoms. Patients with SZ completed fewer years of formal education (*p* = 0.003) and had a higher Body mass index (BMI) (*p* = 0.009) than HC. The SZ group displayed a significantly lower score in PSP as a measure of global level of functioning (*p* < 0.001). Patients with SZ also displayed a lower cognitive score (*p* < 0.001).

**Table 1.**
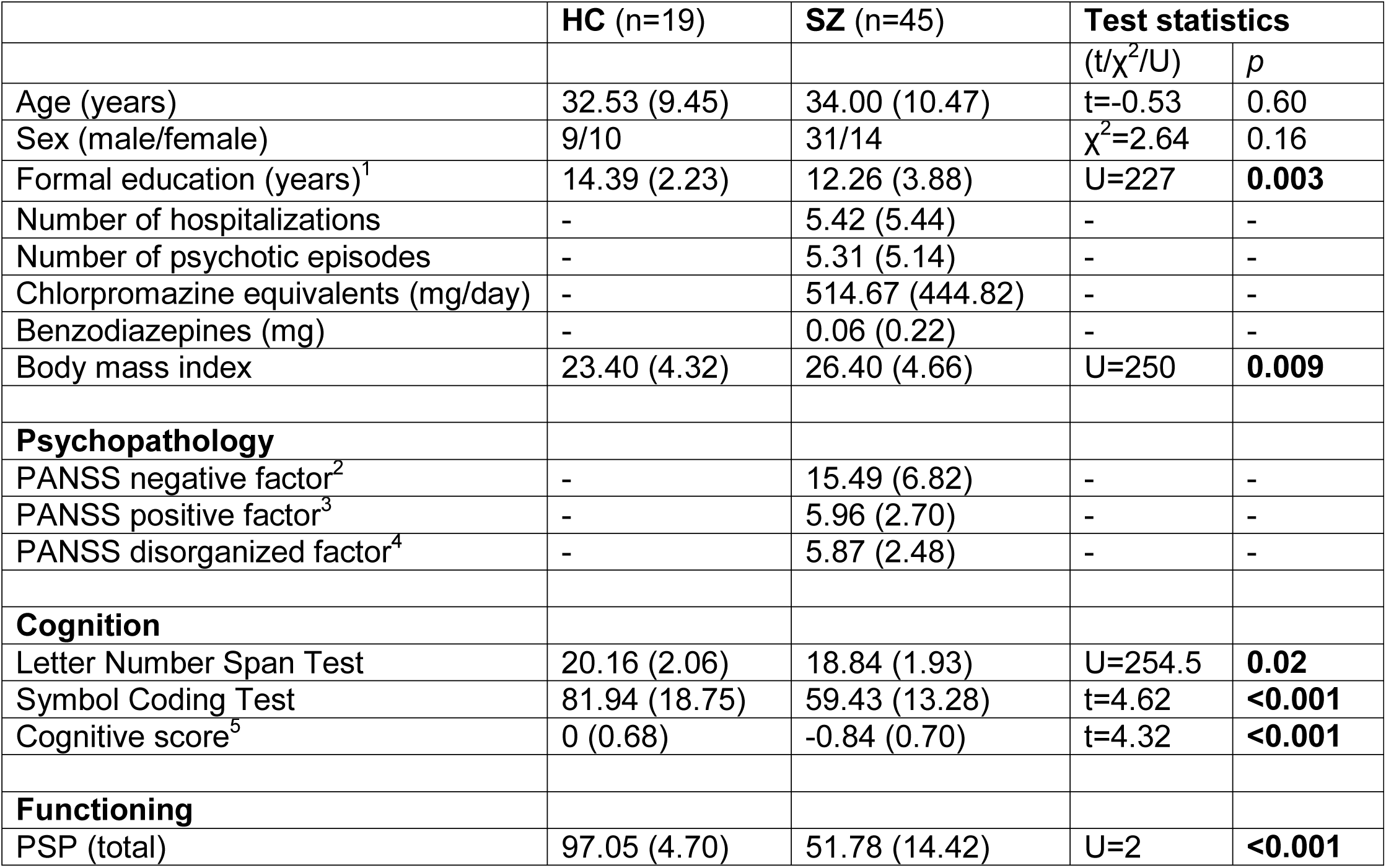
Sample characteristics. Data are presented as means (standard deviations). P values lower than 0.05 are indicated in bold. ^1^Compulsory education in Switzerland is 9 years. ^2^Negative factor: N1, N2, N3, N4, N6, G7. ^3^Positive factor: P1, P3, P5, G9. ^4^Disorganized factor; P2, G5, N11. ^5^Cognition data have been z-transformed based on the data of the healthy control group for each test separately. The cognitive score was computed as the mean of the z-transformed test scores on subject level. Abbreviations: BMI: body mass index; HC: Healthy controls; PANSS: Positive and Negative Syndrome Scale; PSP: Personal and Social Performance Scale; SZ: Schizophrenia.

| **Table 1** | **HC (n=19)** | **SZ (n=45)** | **Test statistics** |  |
| --- | --- | --- | --- | --- |
| | | | (t/$\chi^2$/U) | *p* |
| Age (years) | 32.53 (9.45) | 34.00 (10.47) | t=-0.53 | 0.60 |
| Sex (male/female) | 9/10 | 31/14 | $\chi^2=2.64$ | 0.16 |
| Formal education (years)<sup>1</sup> | 14.39 (2.23) | 12.26 (3.88) | U=227 | **0.003** |
| Number of hospitalizations | - | 5.42 (5.44) | - | - |
| Number of psychotic episodes | - | 5.31 (5.14) | - | - |
| Chlorpromazine equivalents (mg/day) | - | 514.67 (444.82) | - | - |
| Benzodiazepines (mg) | - | 0.06 (0.22) | - | - |
| Body mass index | 23.40 (4.32) | 26.40 (4.66) | U=250 | **0.009** |
| **Psychopathology** |  |  |  |  |
| PANSS negative factor<sup>2</sup> | - | 15.49 (6.82) | - | - |
| PANSS positive factor<sup>3</sup> | - | 5.96 (2.70) | - | - |
| PANSS disorganized factor<sup>4</sup> | - | 5.87 (2.48) | - | - |
| **Cognition** |  |  |  |  |
| Letter Number Span Test | 20.16 (2.06) | 18.84 (1.93) | U=254.5 | **0.02** |
| Symbol Coding Test | 81.94 (18.75) | 59.43 (13.28) | t=4.62 | **<0.001** |
| Cognitive score<sup>5</sup> | 0 (0.68) | -0.84 (0.70) | t=4.32 | **<0.001** |
| **Functioning** |  |  |  |  |
| PSP (total) | 97.05 (4.70) | 51.78 (14.42) | U=2 | **<0.001** |

### 3.2. Patients with SZ display impaired performance in the exploration-exploitation bandit task

First, the total reward earned from the bandit task was calculated for each group. Patients with schizophrenia obtained significantly fewer points than HC (HC: 62.9 ± 5.0; SZ: 59.1 ± 4.7), t = 2.89 *p* = 0.005, d = 0.78) (**Fig. 1A**). Both HC and patients with schizophrenia earned significantly more than the simulated random chooser (random choosers: 51.9, HC: *p* < 0.0001, SZ: *p* < 0.0001). Thus, even though patients with SZ earned less than HC, individuals with SZ clearly performed better than chance.

**Figure 1.**
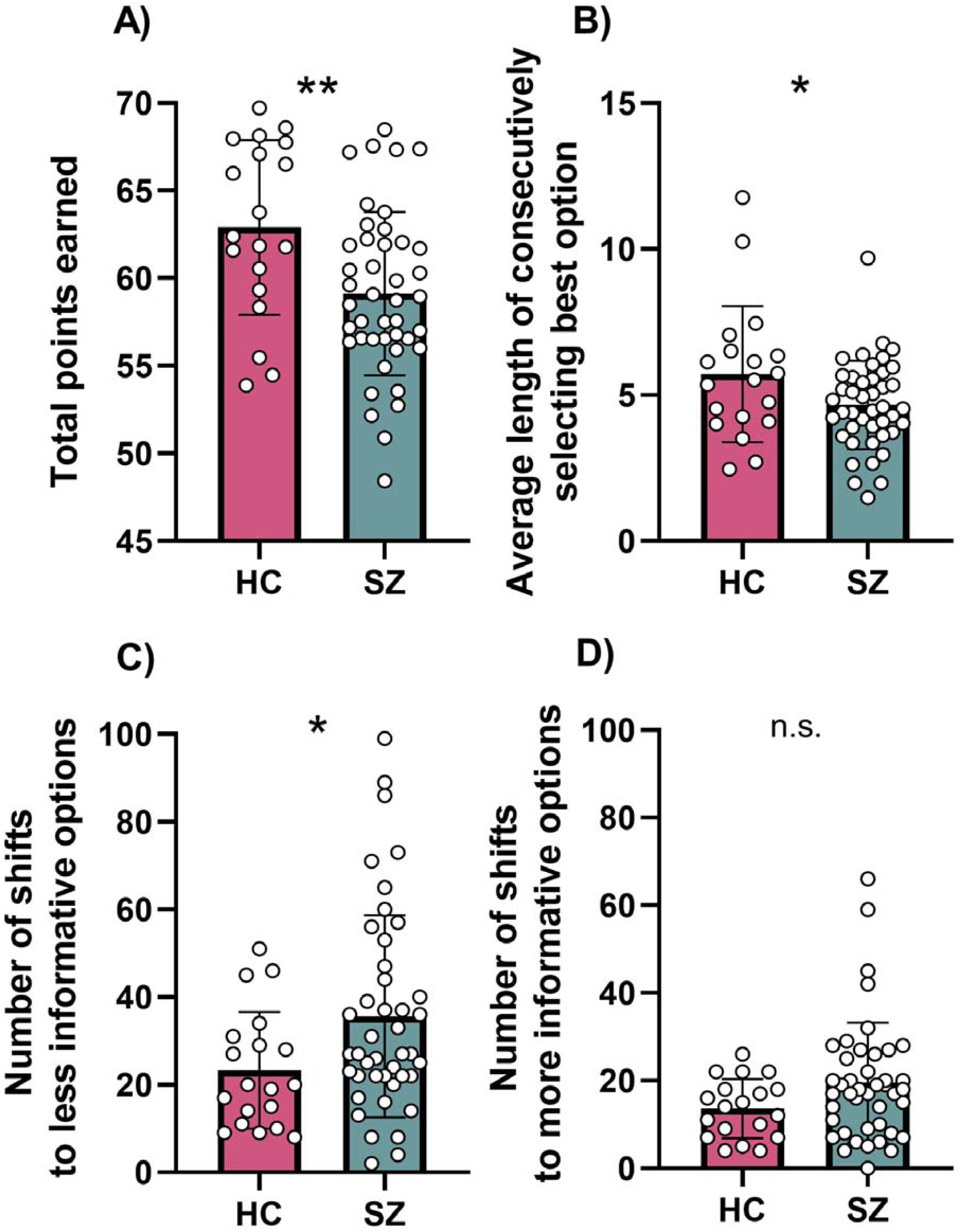
**A)** Patients with schizophrenia (SZ) earned fewer points in a three armed bandit task than healthy controls (HC). Compared to HC, patients with SZ **B)** displayed reduced exploitation of the best option, and showed **C)** more random exploration, but **D)** little differences in directed exploration. * *p* < 0.05, ** *p* < 0.01.

In a next step, we aimed to investigate potential reasons for why patients with SZ earned less reward than HC. To do so, we analyzed exploration and exploitation behaviors. We found that exploitation, defined as the continuous selection of the best bandit option, was reduced in patients with SZ (HC: 5.71 ± 2.3; SZ: 4.65 ± 1.5), t = 2.14, *p* = 0.036, d = 0.55) (**Fig. 1B**). In contrast, there were no significant differences between groups in the length of consecutively selecting other options (*p*’s > 0.7). We then tested whether there were group differences in exploratory behaviors. SZ patients showed more random exploration (HC = 23.32 ± 13.3, SZ = 35.63 ± 23.1; U = 275.5, *p*= 0.042, d = 0.65) **(Fig. 1C)** but similar levels of directed exploration (HC = 13.6 ± 6.7, SZ = 19.6 ± 13.7; U = 302, *p* = 0.104) (**Fig. 1D**). Furthermore, there was a significant negative correlation between random exploration and points earned in the task (r(p) = −0.34, p = 0.025) in patients with SZ.

Next, we performed correlational analyses between the task parameters and three psychopathological domains relevant to schizophrenia: negative symptoms, symptoms of disorganization and cognitive symptoms. Interestingly, symptoms of disorganization, assessed by the PANSS disorganized factor, correlated positively with random exploration (r(s) = 0.31, *p* = 0.045) and negatively with exploitation (r(s) = −0.36, *p* = 0.018), the two task parameters that contributed to the decreased overall task performance in SZ patients compared to HC, but not with directed exploration (r(s) = 0.27, *p* = 0.078). While we observed a significant negative correlation between the PANSS disorganized factor and composite cognitive score (r(s) = −0.33, *p* = 0.029), task parameters did not correlate with cognitive or negative symptoms (**Table S1**). Since antipsychotic medications are known to have an influence on various behaviors and cognition, we correlated chlorpromazine equivalents with task parameters, but did not observe a significant correlation (**Table S1**).

Together, our data show that patients with SZ display impaired task performance since they earned fewer points in the bandit task. Analyzing the underlying behaviors revealed that this deficit is due to an increased random exploration and decreased exploitation, which both correlated with the PANSS disorganized factor.

### 3.3. Patients with SZ show a pro-inflammatory phenotype

To investigate group differences in blood immune profiles between SZ and HC, we used two different exploratory strategies. First, we investigated group differences of individual immune parameters between HC and patients with SZ. Several immune markers differed between groups, indicative of a pro-inflammatory phenotype in patients with SZ compared to HC (**Fig. 2A**): C-reactive protein (CRP) (U = 243.5, *p* = 0.007, d = 0.96), Interleukin (IL)-6 (t = 2.31, *p* = 0.025, d = 0.74), Tumor Necrosis Factor (TNF)-related apoptosis-inducing ligand (TRAIL) (t = 4.16, *p* < 0.001, d = 1.05), TNF-related activation-induced cytokine (TRANCE) (t = 2.09, *p* = 0.041, d = 0.59), Fibroblast Growth Factor 21 (FGF-21) (t = 2.19, *p* = 0.033, d = 0.61), Chemokine Ligand (CCL) 2 (t = 2.05, *p* = 0.045, d = 0.61), CCL7 (t = 2.23, *p* = 0.030, d = 0.71), CCL11 (t = 2.75, *p* = 0.008, d = 0.83) and CCL20 (U = 158.5, *p* < 0.001, d = 1.02) were increased in the SZ group, while Stem Cell Factor (SCF) (t = 2.13, *p* = 0.038, d = 0.62), CXCL11 (U = 223, *p* = 0.026, d = 0.56) and CCL28 (U = 183.5, *p* = 0.003, d = 0.48) were decreased.

**Fig. 2.**
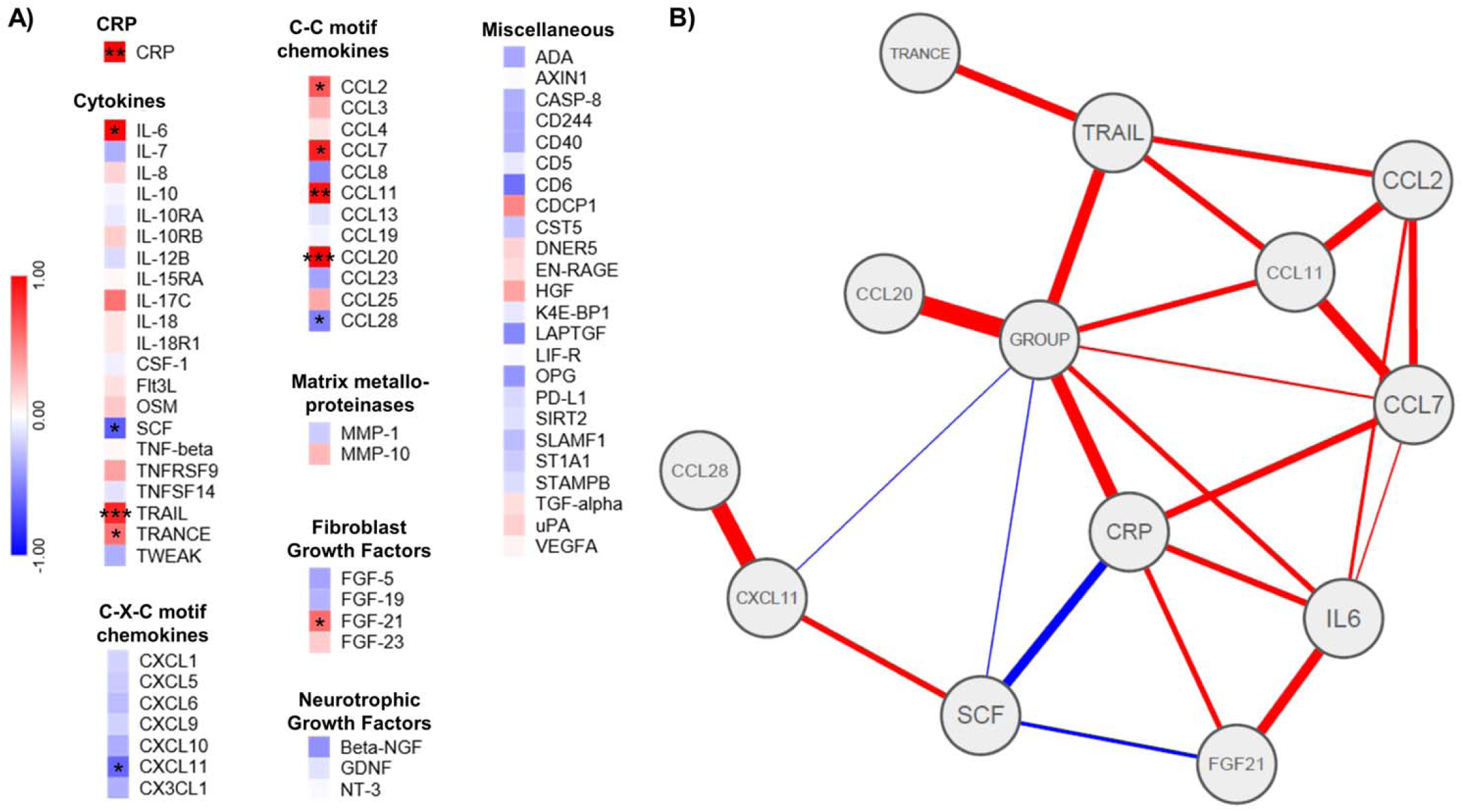
**A)** Heat maps of standardized z-scores of inflammatory markers. Red indicates increased expression and blue decreased expression. **B)** Network analysis. The thickness of an edge corresponds to its weight. Lines indicate positive (red) or negative (blue) partial correlation between two nodes. * p < 0.05, ** p < 0.01, *** p < 0.001. For a list of abbreviations see **Table S2**.

CRP (*p* = 0.022), TRAIL (*p* = 0.001), CCL20 (*p* = 0.003), CCL7 (*p* = 0.027) and CCL11 (*p* = 0.010) remained significantly different after controlling for age, sex and BMI.

We then performed network analysis of the inflammatory markers that were significantly different between the two groups. Given that cytokines and other pro-inflammatory markers are known to influence each other reciprocally, network analysis can reveal distinct associations between group affiliation and the included proteins. Controlling for the associations with all other variables in the network, we found group affiliation to be uniquely correlated with several inflammatory markers, most strongly with CCL20, TRAIL and CRP (**Fig. 2B, Fig. S3** – **S7**).

Together, several pro-inflammatory cytokines and chemokines were elevated in patients with SZ compared to HC, indicative of a pro-inflammatory state. Across all the analyses, CRP, CCL20 and TRAIL were most strongly associated with group.

### 3.4. Random but not directed exploration is associated with increased CRP

Finally, we correlated bandit task behaviors with TRAIL, CCL20 and CRP, the immune parameters that were most strongly associated with group across the analyses. To control for multiple comparisons, significance level was set at *p* < 0.017 (0.05 / 3 parameters).

We found a significant positive correlation of CRP with random exploration (r(s) = 0.40, *p* = 0.008) (**Fig. 3A**), but not exploitation (r(s) = −0.27, *p* = 0.081) (**Fig. 3B**), or directed exploration (r(s) = 0.17, *p* = 0.284) (**Fig. 3C**). The correlation between random exploration and CRP remained significant after controlling for age, BMI, sex, cognition and chlorpromazine equivalents (r = 0.33, *p* = 0.046). There were no correlations of CCL20 or TRAIL with task parameters (**Table S3**).

**Fig. 3.**
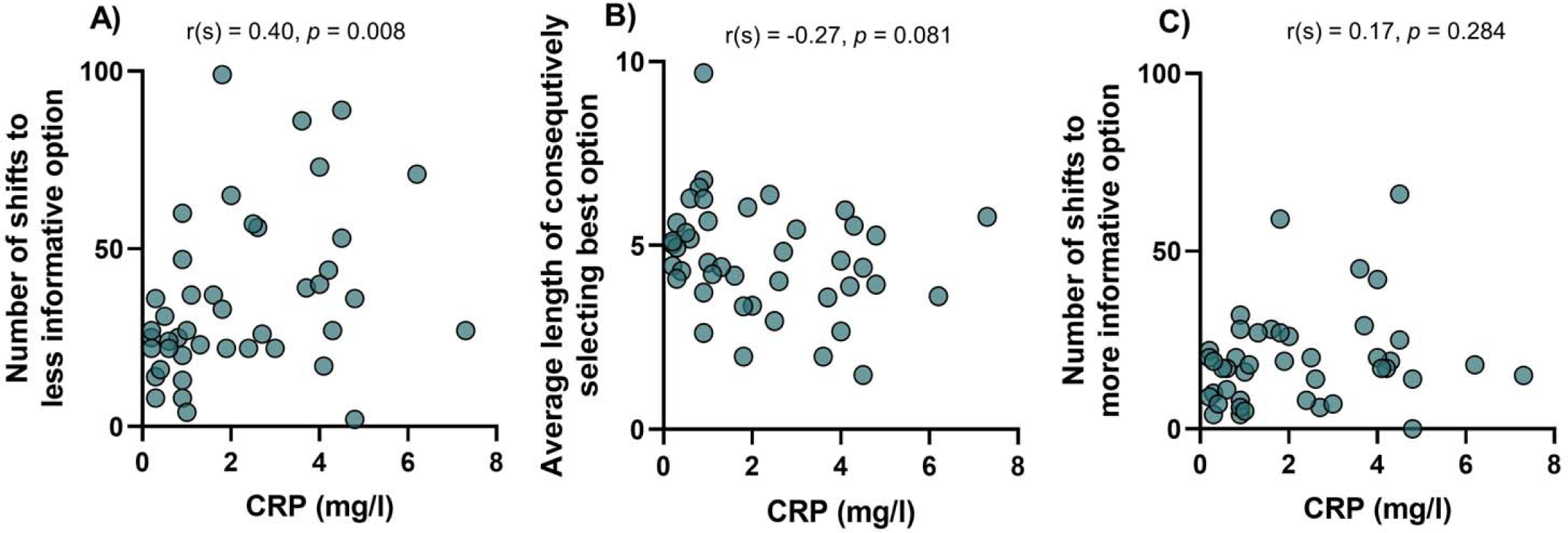
Within patients with schizophrenia, correlation of high sensitivity C-reactive protein (CRP) with **A)** random exploration was significant, in contrast to correlation with **B)** exploitation or **C)** directed exploration.

## Discussion

In the present study, we investigated the link between shifted balancing of exploration versus exploitation using an objective task-based measure, and inflammatory parameters in patients with SZ. Compared to HC, patients with SZ showed reduced performance in the bandit task. Furthermore, we demonstrated that reduced task performance was associated with increased random exploration in SZ. Random exploration correlated with the acute-phase protein CRP.

To our knowledge, the specific increase in random exploration associated with reduced overall performance has not been previously reported for patients with SZ. However, in the study of Martinelli and colleagues, patients with SZ showed increased novelty seeking on a different version of the three-armed bandit task [17]. Although the task and the analysis are not directly comparable, their findings are consistent with ours as novelty seeking and random exploration tap into similar functions. In addition, several studies using simpler tasks have shown that patients with SZ display impaired ability to keep their best decisions and instead more often show “win-shift” behaviors, which is in line with our finding of reduced exploitation [45] [46]. In contrast to findings from our study, Strauss and colleagues showed, using a “temporal utility integration task” that, compared to HC, patients with SZ have decreased uncertainty-driven directed exploration, an effect that was most pronounced in patients with SZ with high levels of negative symptoms [16]. Further, directed exploration was negatively associated with anhedonia in SZ [16]. Although the use of different tasks makes direct comparisons difficult, it is interesting that in the present study patients with SZ did not show differences in directed exploration or correlations of negative symptoms with task parameters.

Instead, we observed a positive correlation between symptoms of disorganization and random exploration. Compared to other symptom domains, symptoms of disorganization have been the least studied, but are closely linked to functional outcomes [47] [2]. Specifically, they have been associated with lack of insight [48], lower quality of life [49] and impaired long-term functioning [50]. In addition, several studies have shown that symptoms of disorganization are related to symptoms of cognition [51] [47]. In line with these findings, we show a negative correlation between PANSS disorganized factor and cognitive score. However, the increased random exploration cannot be solely explained by cognition because we did not find a correlation between cognitive score and task behaviors. Taken together, our data suggest that exploratory behaviors assessed by the bandit task are less relevant to negative symptoms but are more related to symptoms of disorganization, i.e. impairment of integrating complex inputs about a changing environment into an organized, efficient process.

The neural mechanisms underlying exploration-exploitation trade-offs and random exploration in particular have not been fully elucidated yet. Several neurotransmitters, including dopamine, acetylcholine and norepinephrine have been suggested to modulate exploration-exploitation trade-offs [10]. While the neuromodulator dopamine has been consistently implicated in balancing exploration with exploitation [35] [52] [53], the underlying etiological causes of this dopamine dysregulation are not well understood. There is a growing body of evidence indicating that pro-inflammatory cytokines can directly influence dopamine neurotransmission [20]. We therefore investigated if a pro-inflammatory state is associated with the observed task-performance deficits and in extension the hypothesized hypodopaminergic state. In line with previous research [25], patients with SZ showed a pro-inflammatory state. Across all the analyses investigating group differences of inflammatory parameters, CCL20, TRAIL and CRP were most strongly associated with SZ group affiliation. In addition, CRP was positively correlated with random exploration, the task-behavior that contributed to decreased overall task performance.

The acute-phase protein CRP has numerous functions in the body’s immune response, including the induction of cytokine production and activation of the complement pathway [54] [55]. While CRP itself does not cross the blood brain barrier (BBB), peripheral CRP was shown to directly increase the permeability of BBB by reducing tight junction proteins [56]. This increased BBB permeability could facilitate the access of potentially neurotoxic substances such as pro-inflammatory cytokines, peripheral leukocytes or autoantibodies to the brain [25] [57]. Taken together, the current data suggest that patients with SZ show increased pro-inflammatory markers compared to HC and that CRP is associated with increased inefficient, random exploration.

There are several limitations that should be considered when interpreting the results of the current study: First, patients were medicated, and although a recent meta-analysis has not found an effect of antipsychotics on CRP [25], a confounding effect of antipsychotics on both behavior and metabolism cannot be excluded. Moreover, to causally test the hypothesis that peripheral inflammatory mediators can indeed influence neurotransmitter levels leading to behavioral changes, future studies directly measuring dopamine and pharmacologically blocking dopamine receptors in combination with experimentally induced immune challenges in HC, patients with SZ, and pre-clinical models are needed.

Taken together, the present study suggests that patients with SZ are more likely to randomly explore than HC and that this behavior is associated with symptoms of disorganization. Importantly, this inefficient form of exploratory behavior is associated with a pro-inflammatory state. Further studies are needed to elucidate the underlying mechanism of how CRP leads to the neurobiological changes resulting in the observed behavioral alterations. Nevertheless, our data suggest that CRP could be a marker for the severity and treatment course of and potentially a therapeutic target for maladaptive exploratory behaviors in schizophrenia and potentially other neuropsychiatric disorders.

## Acknowledgments

This work was supported by an Advanced Postdoc.Mobility Fellowship of the Swiss National Science Foundation (FC), a Walter and Gertrud Siegenthaler Postdoctoral Fellowship (FC), and project support by the Hartmann Mueller Foundation (FC), Kurt und Senta Herrmann Stiftung (FC) and the Olga Mayenfisch Stiftung (MH). FK was supported by an Early.Postdoc Mobility Fellowship of the Swiss National Science Foundation. PNT acknowledges support by the Swiss NSF (Grants PP00P1 150739 and 100014_165884). TRS was supported by the Forschungskredit of the University of Zurich (FK-19-048).

