## Supplementary Information for "Increased random exploration in schizophrenia is associated with inflammation"

**Figure and Table Legends**

**Fig. S1.** Overview of the bandit task sequence. The three blue squares in every screenshot indicate the individual bandits. After selecting the bandit with the keypad, the points won are displayed in the center of the bandit. RT = Reaction time, ITI: Inter trial interval.

**Fig. S2.** Defining directed and random exploration based on the value of the selected bandit (i.e. random exploration equals switching from the best to the worst and directed exploration switching from the best to the second best option) revealed **A)** increased random (t = 2.31, *p* = 0.019), but not **B)** directed exploration (t = 1.52, *p* = 0.133) of patients with schizophrenia (SZ) compared to healthy control (HC) participants. * *p* < 0.05.

**Fig. S3.** Centrality analysis shows that C-reactive protein (CRP) has the highest strength of all immune-parameters included.

**Fig. S4.** Bootstrap based significance testing of non-zero estimated edge-weights of the network shown in Fig. 2B. Non-significant differences between edge weights are indicated by gray boxes, significant differences by black boxes. Significance is based on α = 0.05. The colour of the diagonal boxes (ranging from red to blue) corresponds to the weights of the edges, with red indicating negative and blue positive weights.

**Fig. S5**. Bootstrap based significance testing of node strength for the nodes of the network shown in Fig. 2B. Non-significant differences between strength of two nodes are indicated by gray boxes, significant differences by black boxes. Node strength of each node is presented by the number in the white boxes.

**Fig. S6.** The average correlation between bootstrap centrality measures of the network estimated with all cases and networks in which some cases were dropped.

**Fig. S7.** Bootstrapped 95% confidence intervals for estimated edge weights of the network shown in Figure 2B.

**Table S1.** Exploitation and random exploration correlate with the Positive and Negative Syndrome Scale (PANSS) disorganized factor but not with PANSS negative factor or cognitive score. There are no associations between task parameters and chlorpromazine equivalents.

**Table S2.** Abbreviations of immune parameters.

**Table S3.** Significant correlation between random exploration with high sensitivity C-reactive protein (CRP), but not with TRAIL or CCL20.

**Supplementary Figures**

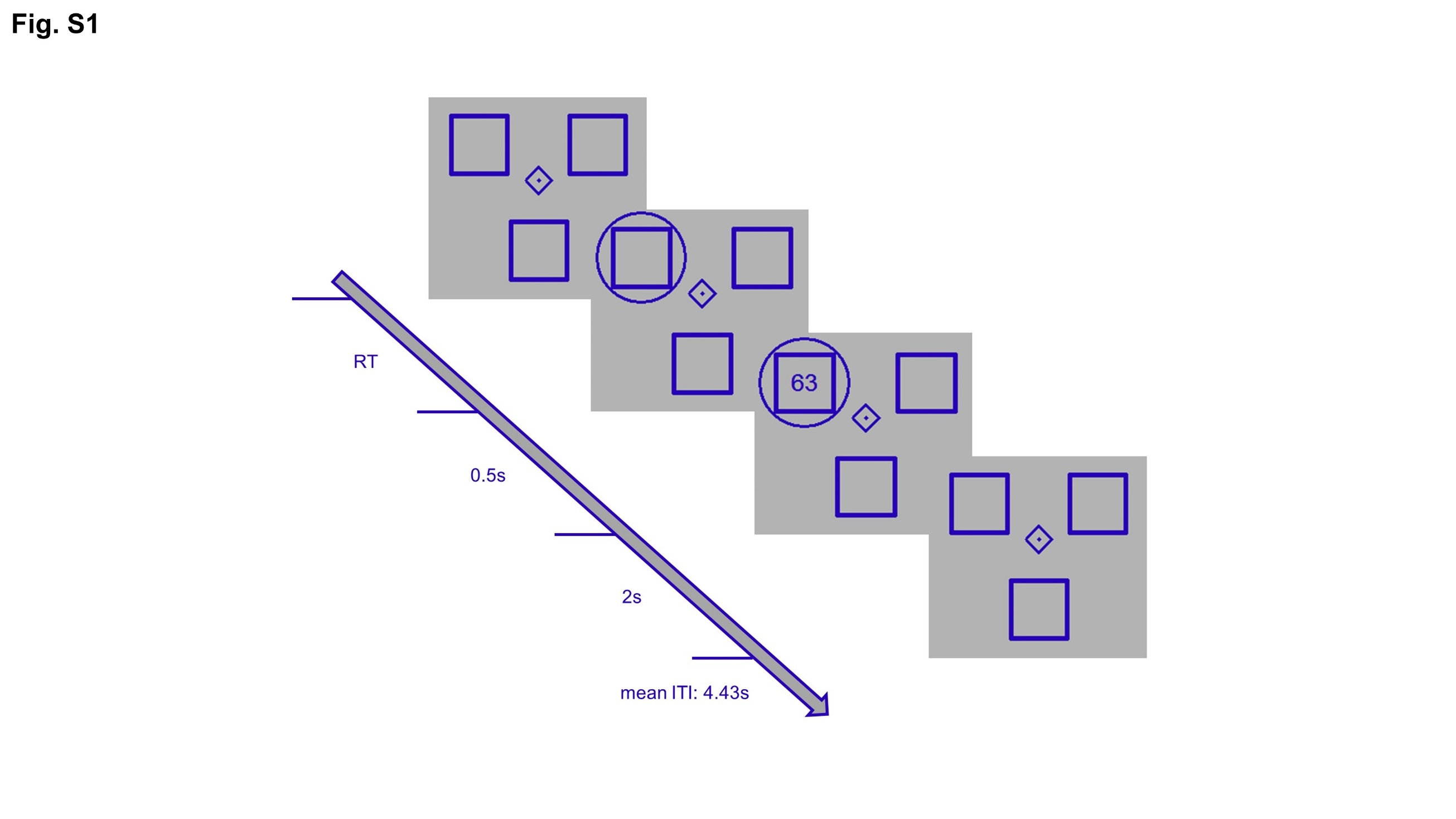
**Figure S1
Figure S2**
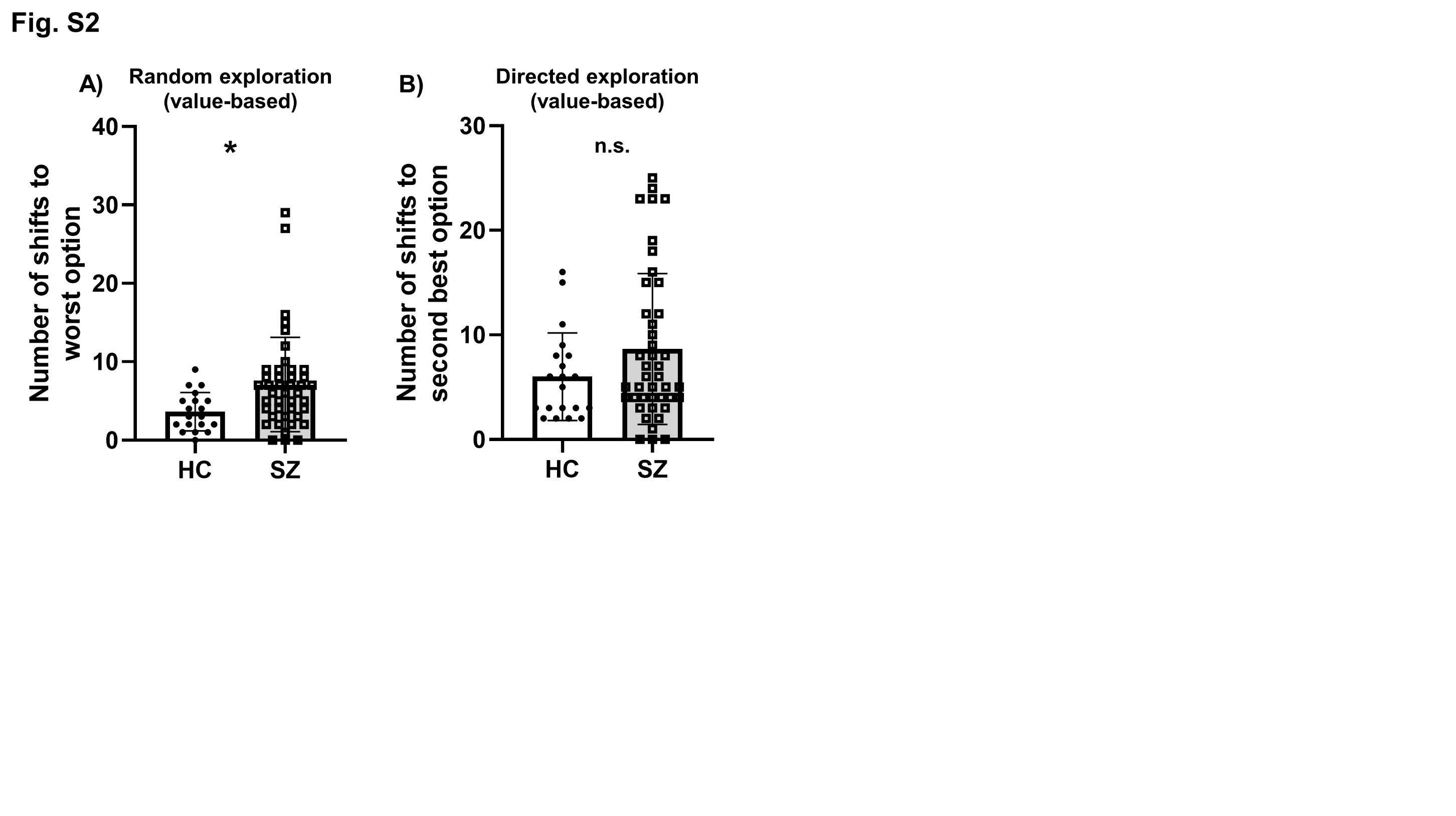

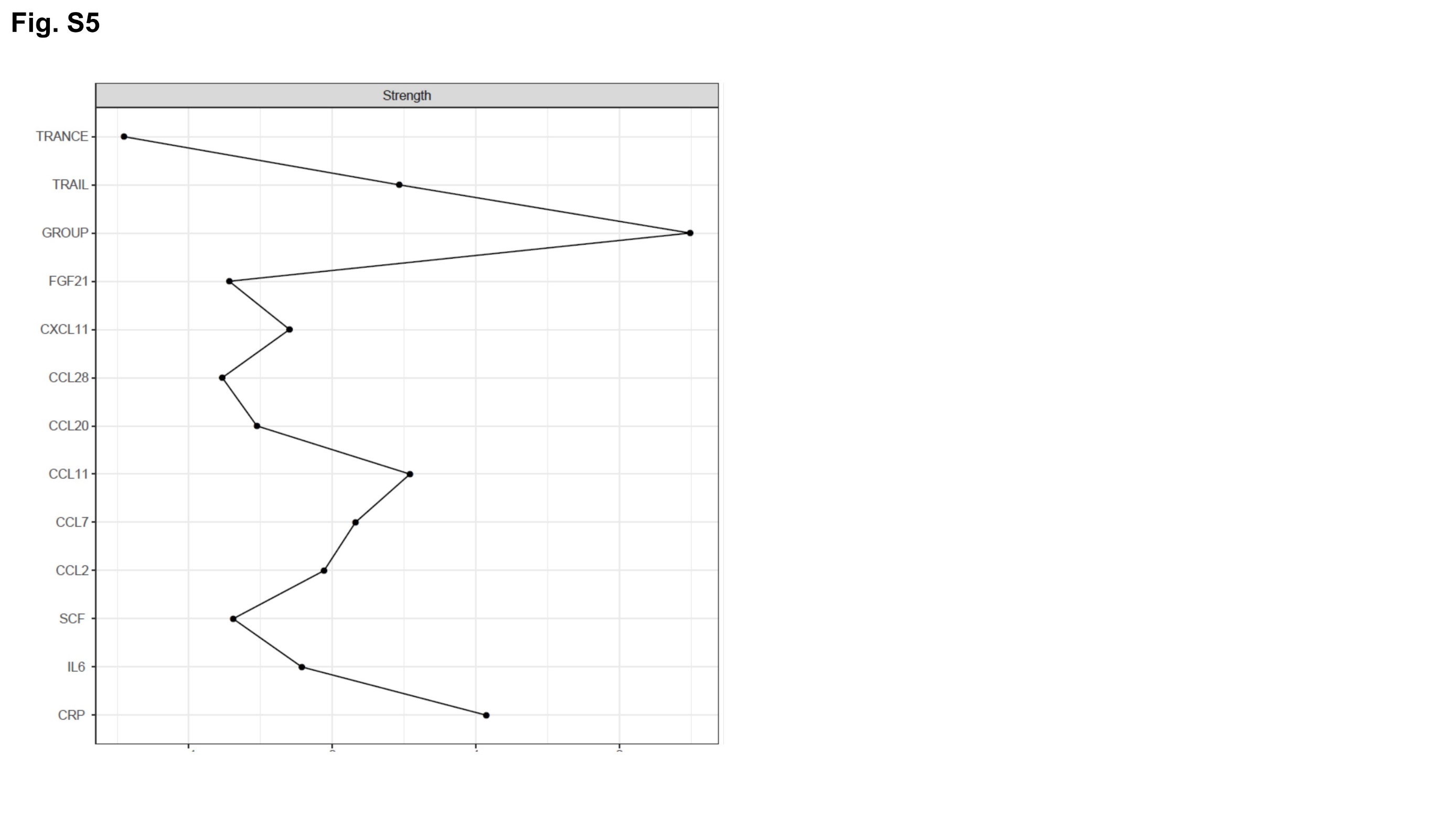
**Figure S3**

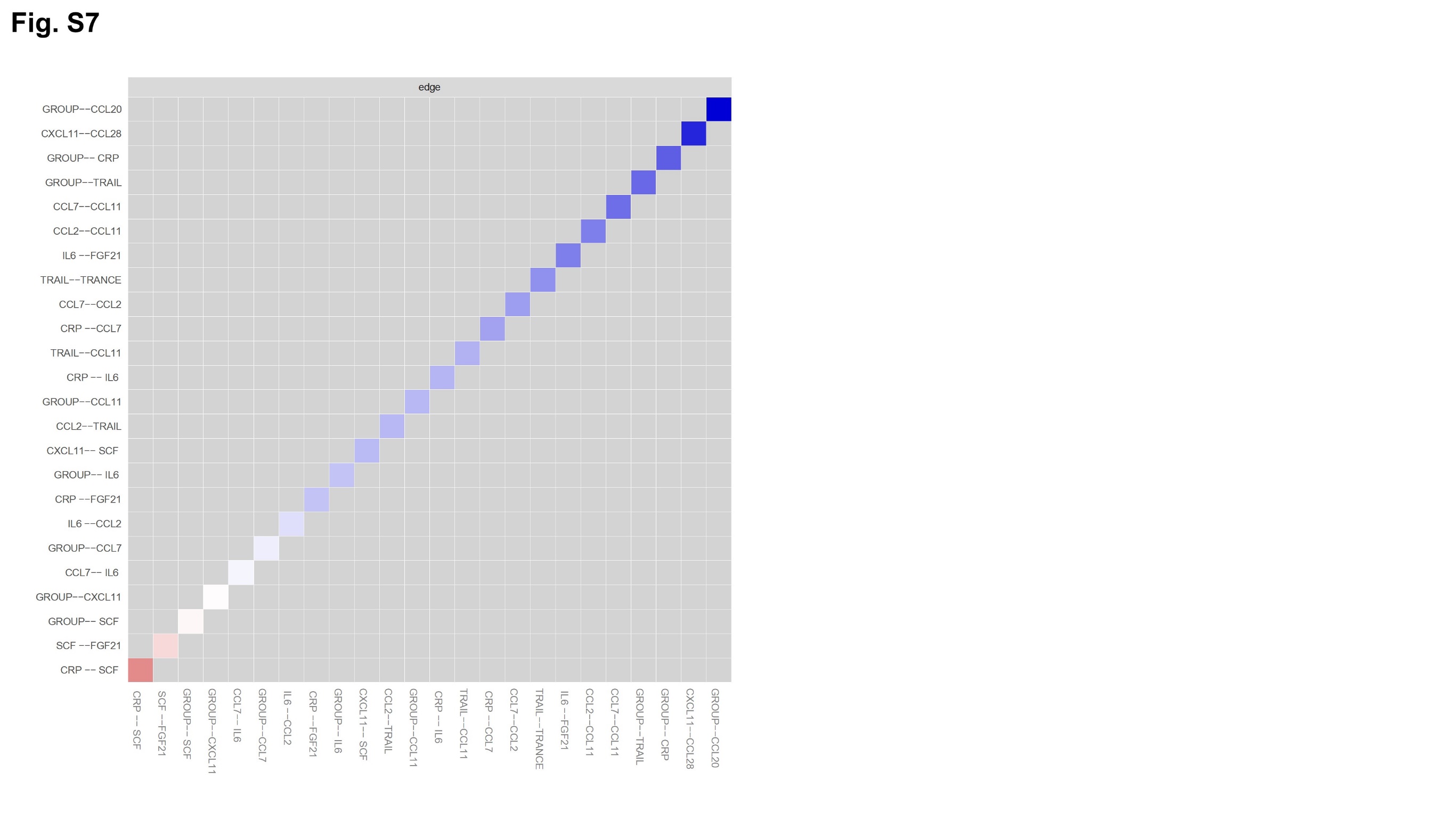
**Figure S4**

**Figure S5**

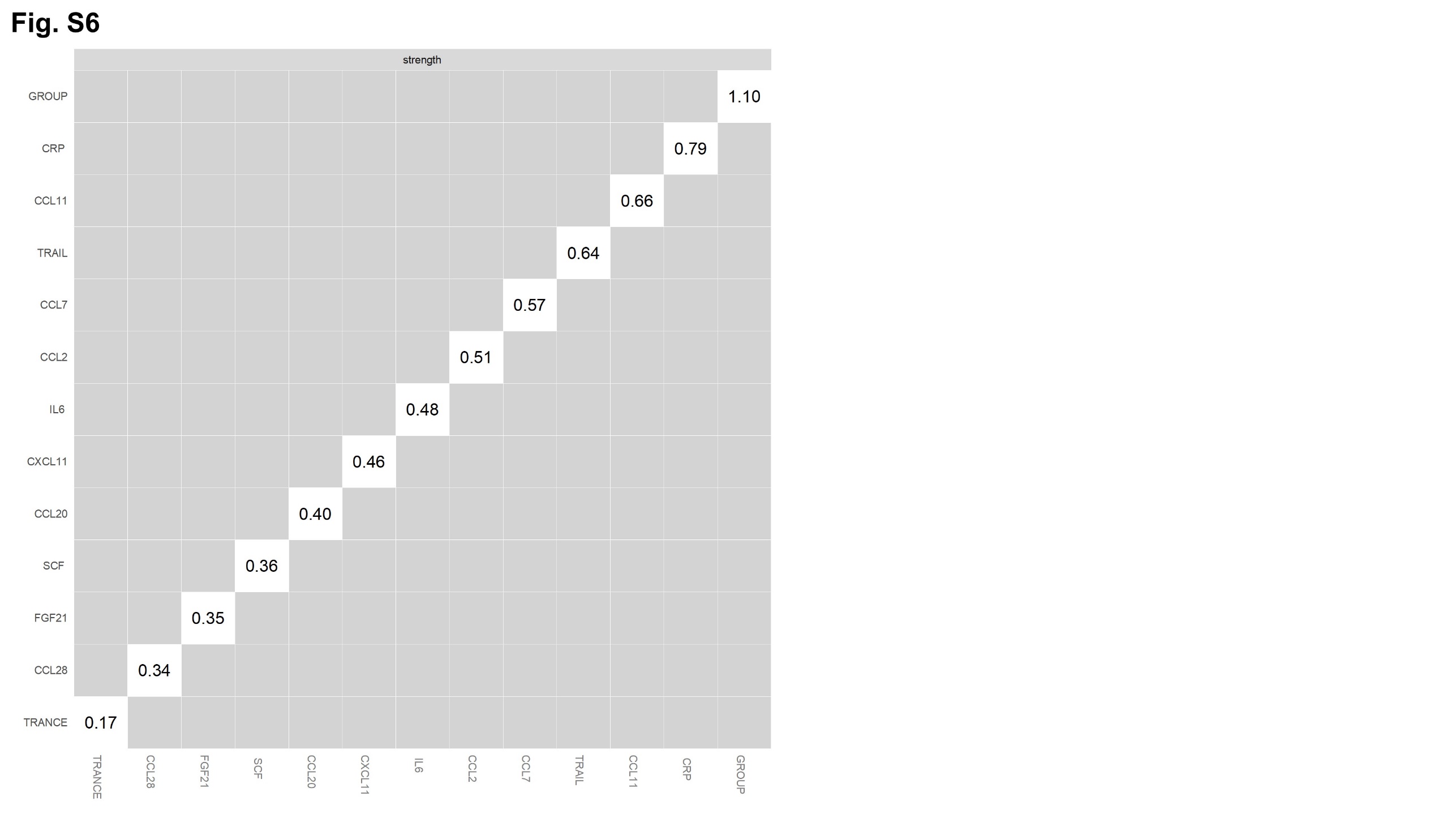

**Figure S6**

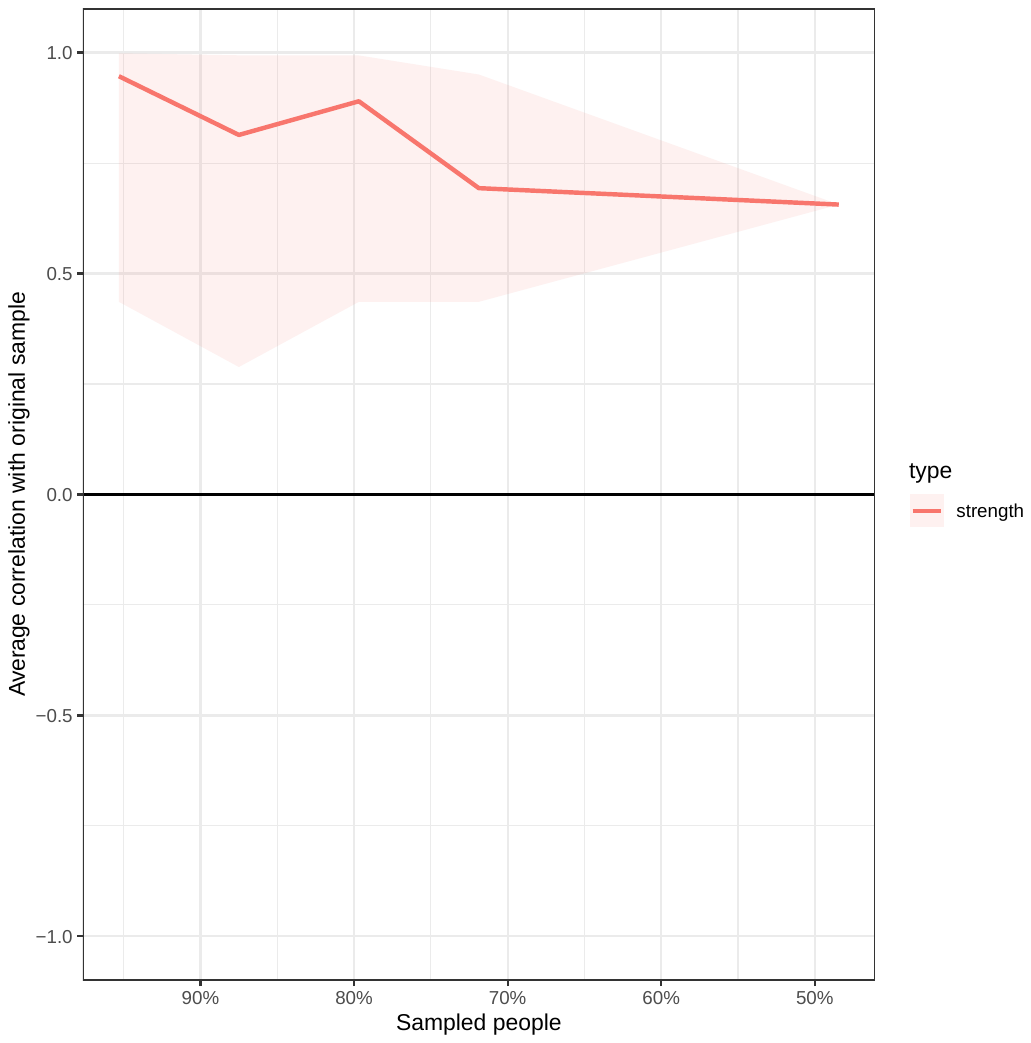

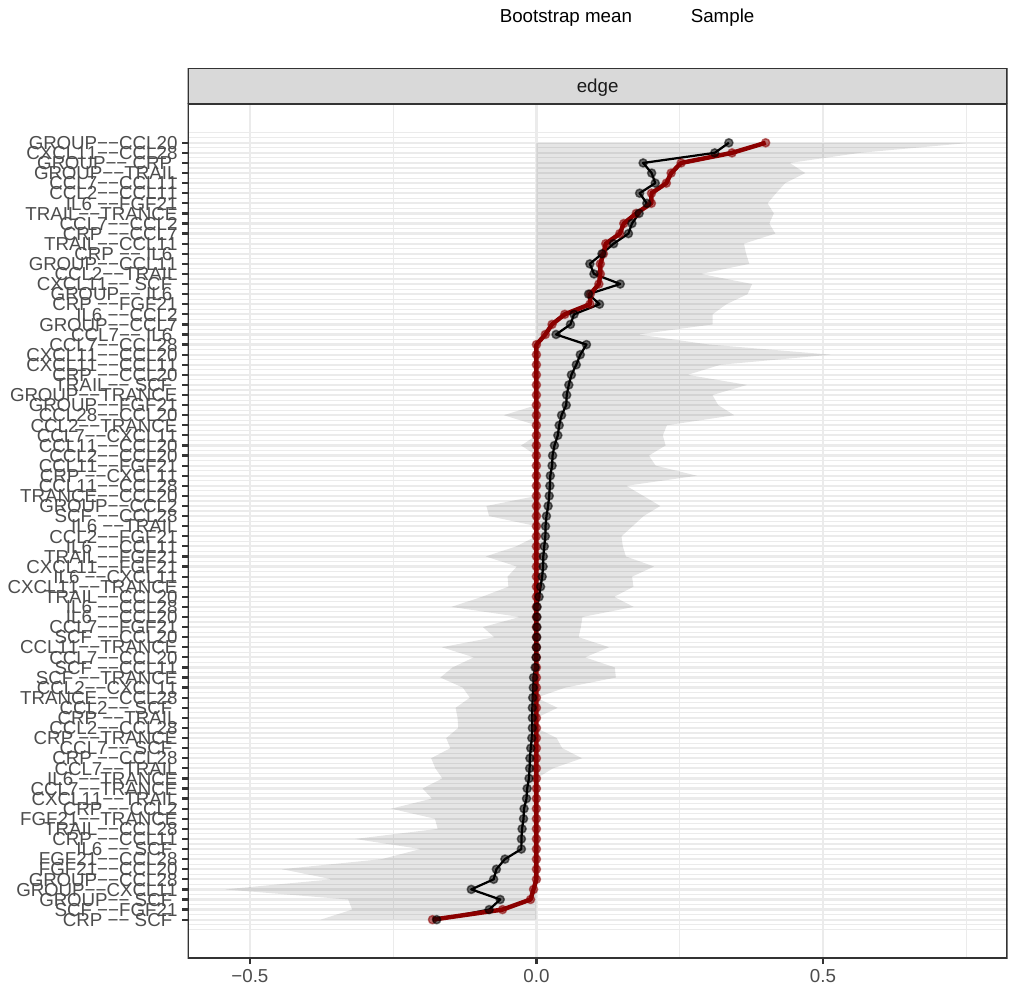
**Figure S7**

**Supplementary Tables**

| **Table S1** | PANSS negative  factor | PANSS disorganized factor | Composite cognitive score | Chlorpro-mazine equivalents |
| --- | --- | --- | --- | --- |
| Total Points  earned | r(s) = - 0.07, *p* = 0.678 | r(s) = -0.30, *p* = 0.055 | r(s) = 0.07, *p* = 0.654 | r(s) = -0.21, *p* = 0.172 |
| Exploitation | r(s) = - 0.09, *p* = 0.558 | **r(s) = -0.36, *p* = 0.018** | r(s) = 0.16, *p* = 0.303 | r(s) = -0.22, *p* = 0.159 |
| Random  exploration | r(s) = 0.14, *p* = 0.375 | **r(s) = 0.31, *p* = 0.045** | r(s) = -0.18, *p* = 0.454 | r(s) = 0.23, *p* = 0.132 |
| Directed exploration | r(s) = -0.01, *p* = 0.967 | r(s) = 0.27, *p* = 0.078 | r(s) = 0.01, *p* = 0.935 | r(s) = 0.15, *p* = 0.338 |

**Correlations between task-parameters and psychopathology, composite**

**cognitive score and chlorpromazine equivalents in patients with SZ**

| **Table S2** |  |
| --- | --- |
| **Abbreviation** | **Legend** |
| ADA | Adenosine Deaminase |
| AXIN1 | Axin-1 |
| Beta-NGF | Beta-nerve growth factor |
| CASP-8 | Caspase-8 |
| CCL | C-C motif chemokine |
| CD | Cluster of differentiation |
| CDCP1 | CUB domain-containing protein 1 |
| CRP | C-reactive protein |
| CSF-1 | Colony-stimulating factor 1 |
| CST5 | Cystatin-D |
| CXCL | C-X-C motif chemokine |
| DNER5 | Delta and notch-like epidermal growth factor-related receptor |
| EN-RAGE | Protein S100-A12 |
| Flt3L | Fms-related tyrosine kinase 3 ligand |
| FGF | Fibroblast growth factor |
| GDNF | Glial cell line-derived neurotrophic factor |
| HGF | Hepatocyte growth factor |
| IL | Interleukin |
| IL-R | Interleukin receptor |
| LAPTGF | Latency-associated peptide transforming growth factor beta-1 |
| Lif-R | Leukemia inhibitory factor receptor |
| MMP | Matrix metalloproteinase |
| NT-3 | Neurotrophin-3 |
| OPG | Osteoprotegerin |
| OSM | Oncostatin-M |
| PD-L1 | Programmed cell death 1 ligand 1 |
| SCF | Stem cell factor |
| TNFB | Tumor necrosis factor beta |
| TNFRSF9 | Tumor necrosis factor receptor superfamily member 9 |
| TNFSF14 | Tumor necrosis factor ligand superfamily member 14 |
| TRANCE | Tumor necrosis factor-related activation-induced cytokine |
| TWEAK | Tumor necrosis factor (ligand) superfamily, member 12 |
| TRAIL | Tumor necrosis factor related apoptosis-inducing ligand |
| SLAMF1 | Signaling lymphocytic activation molecule family member 1 |
| SIRT2 | SIR2-like protein 2 |
| ST1A1 | Sulfotransferase 1A1 |
| STAMPB | STAM-binding protein |
| uPA | Urokinase-type plasminogen activator |
| VEGFA | Vascular endothelial growth factor |

**List of abbreviations of inflammatory parameters**

| **Table S3** | Total points earned (Reward) | Continuously best (N)  (Exploitation) | Shifts to less informative option (N) (Random Exploration) | Shifts to more informative option (N) (Directed Exploration) |
| --- | --- | --- | --- | --- |
| CRP | r(s) = -0.19, *p* = 0.225 | r(s) = -0.27, *p* = 0.081 | **r(s) = 0.40, *p* = 0.008** | r(s) = 0.17, *p* = 0.284 |
| TRAIL | r(p) = -0.11, *p* = 0.517 | r(p) = -0.12, *p* = 0.445 | r(s) = 0.19, *p* = 0.248 | r(s) = 0.16, *p* = 0.326 |
| CCL20 | r(s) = -0.21, *p* = 0.196 | r(s) = 0.06, *p* = 0.737 | r(s) = 0.10, *p* = 0.557 | r(s) = 0.13, *p* = 0.414 |

**Correlations between CRP, TRAIL and CCL20 with bandit task parameters in patients with SZ**
